# How the non-motile kinesin KIF7 adapts conserved kinesin principles for its function in Hedgehog signaling

**DOI:** 10.64898/2026.02.06.704517

**Authors:** Farah Haque, Bindu Y. Srinivasu, John R. Engen, Thomas E. Wales, Radhika Subramanian

## Abstract

KIF7 is an atypical, non-motile kinesin that regulates Hedgehog signaling by concentrating GLI transcription factors at the cilium tip. How canonical kinesin principles for intracellular transport are repurposed to support KIF7’s function as a signaling scaffold remains unclear. KIF7 exists in an autoinhibited state that is relieved by GLI binding, promoting microtubule association. We examined this regulatory mechanism by combining HDX-MS and AlphaFold modeling of a minimal KIF7 dimer, both alone and in complex with the GLI2 zinc-finger domain. Our HDX-MS data indicate that the highly negatively charged neck-coil dimerization domain of KIF7, which serves as the GLI2-binding site, is intramolecularly protected in the absence of GLI2. Consistent with this, AlphaFold models suggest that the motor domain folds back onto the neck-coil via KIF7’s unusually long neck-linker, sterically occluding the microtubule-binding interface. This occurs through a mechanism conceptually analogous to, but structurally distinct from, autoinhibition in motile kinesins. GLI2 binding to the KIF7 neck-coil displaces the motor domain and induces allosteric changes that propagate to the microtubule-binding surface, thereby activating microtubule binding. ATP turnover further modulates KIF7’s microtubule binding–unbinding equilibrium. Together, these findings reveal how a kinesin is adapted for a non-motile function as a scaffold in Hedgehog signaling.

## Introduction

Kinesins are a large family of microtubule-associated motor proteins that play essential roles in intracellular cargo transport. Most kinesins are motile, using the energy derived from ATP hydrolysis to step directionally along microtubules. This motility is tightly regulated, and one of the best-established paradigms of kinesin regulation is autoinhibition. In the absence of cargo, autoinhibition suppresses the kinesin–microtubule interaction, thereby preventing futile ATP hydrolysis and microtubule crowding^1^. Association with cargoes or cargo adaptors relieves this inhibition, enabling kinesins to bind microtubules and initiate motility.

In contrast to motile kinesins, the kinesin-4 family protein KIF7 is an atypical kinesin that binds microtubules but lacks motility. KIF7 is a conserved regulator of the vertebrate Hedgehog (Hh) signaling pathway that requires the primary cilium^2,3^. Upon pathway activation, KIF7 is transported by the intraflagellar transport (IFT) machinery to the distal tip of the cilium^4^. KIF7 directly binds Glioma-associated oncogene (GLI) transcription factors, the primary effectors of Hh signaling, and concentrates them at the cilium tip for their proper activation^5–8^. Thus, KIF7 functions effectively as a non-motor microtubule-associated protein, despite retaining a canonical kinesin fold and an intact ATPase pocket. This raises a fundamental question: why has Hedgehog signaling co-opted a kinesin, rather than a non-motor MAP (microtubule-associated protein), for this function? Although prior work has identified mechanisms that disrupt mechanochemical coupling and motility in KIF7, how conserved kinesin features are repurposed to support its signaling function remains poorly understood.

The domain architecture of KIF7 shows both conserved and distinct features relative to other kinesins **(Figure 1)**. At the N-terminus is the motor domain (KIF7-motor), which adopts the canonical kinesin fold and possesses ATPase activity^9^. However, unlike typical motile kinesins, where microtubules stimulate basal ATPase rates by ∼1000-fold, KIF7 shows only a modest 3–4-fold increase in ATPase activity in the presence of microtubules^10^. The functional significance of this weak microtubule-stimulated ATPase activity remains unknown. In addition, in KIF7, the microtubule binding–unbinding cycle is not tightly coupled to the nucleotide state, as observed for motile kinesins^11^.

**Figure 1.**
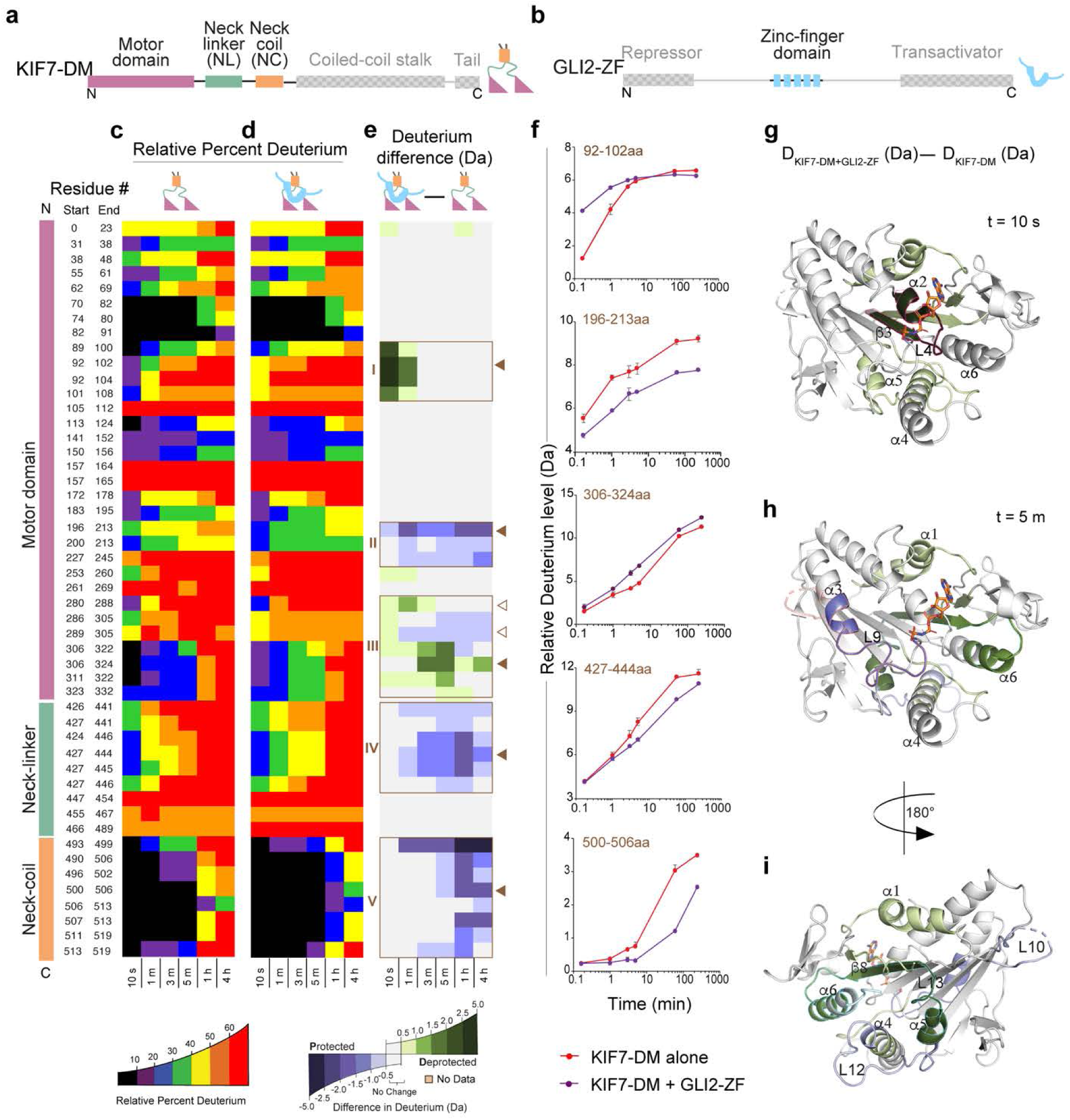
**HDX-MS reveals long-range changes in the structural dynamics of KIF7-DM upon GLI binding** (a) Domain organization of KIF7. Domains present in KIF7-DM: motor domain (pink), neck-linker (green), and neck-coil (orange), and domains not present in KIF7-DM (grey). (b) Domain organization of GLI2. GLI2-ZF domain is in blue, and the sequences not present in GLI2-ZF are in grey. (c) HDX-MS data for curated peptides in KIF7-DM. A Chiclet plot of the relative percentage of deuterium for KIF7-DM is shown. The X-axis shows the time points, and the Y-axis shows each peptide. The relative percentage of deuterium is colored according to the scale shown. (d) Chiclet plot of relative percentage deuterium for the KIF7-DM:GLI2-ZF complex. The X-axis shows the time points, and the Y-axis shows each peptide. The relative percentage of deuterium is colored according to the scale shown. (e) Deuterium difference between (c) and (d). The data are calculated by subtracting the absolute deuterium incorporation of KIF7-DM from the KIF7-DM:GLI2-ZF complex. The X-axis shows the time points, and the Y-axis shows each peptide. Only the peptides that are coincident in both KIF7-DM and KIF7-DM:GLI2-ZF are shown. The differences are colored according to the scale shown. (f) Deuterium incorporation curves of selected peptides (indicated with solid brown arrowheads) in (c) are shown. (g) Deuterium difference of the selected peptides from 10 s time point in (e) are mapped on KIF7 motor domain (PDB ID: <u>7RX0</u>) and colored according to the deuterium difference color scale. (h) and (i) Deuterium difference of the selected peptides from 5 m time point in (e) are mapped on KIF7 motor domain (PDB ID: <u>7RX0</u>) and colored according to the deuterium difference color scale.

Immediately following the motor domain is the neck-linker (KIF7-NL), another conserved element of kinesins. Most kinesins contain a relatively short neck-linker (∼14–18 residues) that enables tight coordination between motor domains during processive stepping^12,13^. In contrast, KIF7 has an unusually long (∼120-residue) neck-linker, predicted to be largely unstructured except for a short α-helical segment (KIF7-NL_helix_), whose functional significance remains unclear. The KIF7-NL is followed by the first coiled-coil domain (∼100 amino acid residues), which is essential for dimerization and is referred to as the neck-coil in kinesins (KIF7-NC)^1^. In KIF7, this domain is characterized by a highly negative surface charge potential^3^. The remainder of the molecule comprises the stalk domain (KIF7-stalk), which contains additional coiled-coil regions and linker sequences, and a C-terminal unstructured tail domain (KIF7-tail). In motile kinesins, the tail domain typically mediates binding to cargoes or cargo adaptors.

Unlike motile kinesins, KIF7 does not bind its known “cargo”, the GLI transcription factors, through its C-terminal tail domain. Instead, the negatively charged KIF7-NC interacts with the DNA-binding zinc-finger domain of GLI2 (GLI2-ZF; 418–594 aa of isoform 2, the primary activator isoform) through principles of DNA mimicry^3^. In addition, GLI2-ZF also binds the KIF7 motor domain^3^. Notably, both of these GLI-binding sites are present in the N-terminal region of KIF7 (KIF7-DM; 1-543aa). Biochemical assays using recombinant proteins show that GLI2-ZF enhances the microtubule affinity of KIF7-DM^3^. These findings indicate that KIF7-DM exists in equilibrium between an autoinhibited and a microtubule-binding-competent state, with GLI2-ZF binding shifting the equilibrium toward the active state. In cells, KIF7’s binding to cytoplasmic microtubules and its localization to the cilium tip following pathway activation depend on its interaction with GLI proteins^3^. Notably, full-length KIF7 is even more stringently autoinhibited than the minimal KIF7-DM, likely through additional contacts involving the C-terminal tail^2^. Together, these observations suggest a conventional kinesin–cargo relationship between KIF7 and GLI, in which GLI binding shifts KIF7 from an autoinhibited to an active state.

At present, the structural basis of this autoinhibited-to-active transition in KIF7, and how it compares with autoinhibition–activation paradigms established for motile kinesins, remains unresolved. Experimentally determined structures of the autoinhibited state are not yet available, and cryo-EM analysis of the microtubule-bound KIF7-GLI2 complex resolved only the motor domain, leaving the neck-linker (KIF7-NL) and the GLI2-bound neck-coil (KIF7-NC) unresolved, likely due to conformational flexibility^3^. Hydrogen–deuterium exchange mass spectrometry (HDX-MS) provides a powerful approach to probe such dynamic, multi-domain rearrangements^14,15^, yet has not been extensively applied to dissect the structural mechanisms of KIF7 regulation. Here, we combine HDX-MS with AlphaFold modeling to investigate the structural mechanisms that regulate KIF7 autoinhibition and its activation by GLI2. Our findings reveal how conserved kinesin regulatory principles are repurposed for a motility-independent function by a non-motile kinesin during Hedgehog signaling.

## Results

### HDX-MS reveals allosteric conformational changes in KIF7-DM upon GLI2-ZF binding

To examine conformational changes in KIF7 upon GLI2 binding, we performed differential HDX-MS experiments. HDX-MS exploits the exchange of backbone amide hydrogens with deuterium, providing a sensitive measure of local solvent accessibility and conformational flexibility^16^. Solvent-exposed or intrinsically dynamic regions typically undergo rapid exchange with deuterium, whereas buried, hydrogen-bonded, or interaction-stabilized segments exhibit markedly slower deuterium uptake^17^. Since full-length recombinant KIF7 and GLI2 are challenging to purify, we used a previously well-characterized recombinant minimal KIF7 dimer (1-543aa; KIF7-DM)^3^ that contains both the GLI2 and microtubule interaction sites **(Figure 1a)** for HDX-MS experiments. Prior studies have shown that the microtubule binding of KIF7-DM is enhanced upon binding to the GLI2 zinc-finger domain (418-594aa; GLI2-ZF)^3^ **(Figure 1b)**. Thus, KIF7DM and GLI2-ZF provide a biochemically tractable system for examining the mechanism of KIF7 autoinhibition and its activation upon GLI2 binding.

We first performed HDX-MS of KIF7-DM in the absence of GLI2-ZF at different time points: 10 s, 30 s, 1 m, 3 m, 5 m, and 1 h of D_2_O exposure. After each specific exposure time the deuterium labeling reactions were quenched by lowering the pH and temperature, digested online using Nepenthesin II, and the levels of deuterium incorporation were analyzed for each peptide by mass spectrometry. The analysis yielded 82.9% KIF7-DM sequence coverage with 2.56 redundancy, indicating robust peptide representation across the protein **(All HDX-MS data are shown in the Supplementary Data File Tabs 1-3)**. The HDX profile revealed variable backbone dynamics across the KIF7-DM sequence. Peptides in two regions showed the lowest HDX (i.e., the lowest relative deuterium incorporation), particularly at the earliest labeling time point (10 s) **(Figure 1c).** The first, as expected, corresponded to the structural core of the motor domain (α1-helix, β3-strand, and P-loop; 70-100aa). Notably, the second corresponded to the neck-coil dimerization domain (KIF7-NC; 493-519aa). The low deuterium incorporation in both regions persisted for 5 min of labeling and, for some peptides, up to 1 h, consistent with a stable structure and slow backbone dynamics. Additionally, the raw MS data for peptides in KIF7-NC showed non-EX2-like behavior^18^, indicating the presence of at least two different exchanging populations in solution **(Supplementary Data File, Tab non-EX2)**.

In contrast to the motor core and the KIF7-NC, the neck-linker (KIF7-NL) and several loops within the motor domain showed higher HDX, as expected for predicted unstructured and flexible regions. In particular, the C-terminus of the KIF7-NL (445-489aa) showed near maximum deuterium incorporation at all labeling time points, indicating high solvent exposure and minimal hydrogen-bonding. Multiple loops in the KIF7 motor domain [L5 (105-112aa), L8 (157-165aa), L10 (227-245aa), and L11 (261-269aa)] showed similarly high HDX. Other regions in the motor domain showed intermediate HDX, including the switch I region (195-213aa) and the microtubule-binding interface (280-332aa). Within this subset, α5-helix, loop L12, and β8-strand (306-332aa) stood out as a region with low HDX even at the earliest experimental time point (10 s), indicating that some areas of the KIF7 microtubule-binding interface are characterized by slower backbone dynamics even in the absence of microtubules.

We next performed differential HDX-MS on the recombinant size-exclusion chromatography (SEC)-purified KIF7-DM:GLI2-ZF complex **(Extended Data Figure 1a-b)** and compared it with KIF7-DM alone. Before initiating the HDX-MS experiments, the integrity of the complex was verified by intact mass analysis **(Supplementary Data File Tab, Intact Mass Analysis)**. The HDX profile of KIF7-DM:GLI2-ZF showed extensive changes compared to KIF7-DM in all domains (NC, NL, and motor) **(Figure 1d)**. First, within the KIF7-NC domain (493-519aa), we observed decreased HDX at later time points in the complex relative to KIF7-DM alone **(>1 hr, Figure 1e V, 1f)**. These changes were not restricted to amino acids previously shown to be important for this protein-protein interaction^3^. Instead, the decrease in HDX was observed throughout the neck-coil region. In addition, the multiple populations observed in the neck-coil of KIF7-DM collapsed into a single population in the KIF7-DM:GLI2-ZF complex (**Supplementary Data File, Tab non-EX2**). These findings suggest a further reduction in structural dynamics and increased stability of the KIF7-NC domain when it is in complex with GLI2-ZF **(Supplementary Data File, Tab non-EX2)**. Next, within the KIF7-NL, a decrease in HDX was observed in the KIF7-NL_helix_ (426–446 aa) in the complex compared with KIF7-DM alone **(Figure 1e IV, 1f)**. These changes in the neck-linker helix were unexpected, given that this domain does not directly bind GLI2-ZF^3^.

Finally, in the KIF7 motor domain, we detected both increases and decreases in HDX in localized regions of the complex relative to KIF7-DM. Notably, the changes were observed in regions of the motor domain distal to the known interaction site of GLI2-ZF on the KIF7 motor domain^3^ **(Extended Data Figure 1d)**. Two regions of the complex showed significant changes in HDX relative to KIF7-DM: the microtubule-binding interface and the ATPase pocket. Previous Cryo EM structures and mutagenesis studies showed that the KIF7-microtubule binding interface is formed by helices α4, α5, and α6, and the loop L12 **(Figure 1e III, 1f, 1g-i)**. We observed: 1) increased HDX in α4-helix (280-288aa), 2) decreased HDX in the positively charged K-loop or Loop L12 (289-305aa), and 3) increased HDX in α5 and α6 helices (303-322aa and 323-332aa). In the ATPase pocket, we observed: 1) protection at all time points in the switch I region (227-245aa), which is critical for the coupling between microtubule binding and ATPase cycle in kinesin^19^ **(Figure 1e II, 1f, g-i),** and 2) increased HDX in the P-loop which contains the nucleotide binding cleft (92-104 aa) at early time points **(<1 min, Figure 1e I, 1f, 1g-i)**. Because these HDX changes map to both the microtubule-binding interface and ATPase pocket, we next examined whether GLI2-ZF binding alters ATPase activity and found a three-fold increase in the microtubule-stimulated ATPase rate of KIF7 in the presence of GLI2-ZF **(Extended Data Figure 1c)**.

Collectively, these findings define the HDX profile of KIF7-DM and how it changes upon GLI2-ZF binding. The data reveal that GLI2-ZF–induced changes extend to regions of KIF7-DM outside of known GLI2-ZF interaction sites, including the neck-linker, the microtubule-binding region, and the ATPase pocket of the motor domain. Together, these observations indicate that GLI2-ZF binding induces significant allosteric conformational changes in KIF7-DM.

### Structural dynamics at the KIF7 motor domain are linked to changes in its neck-coil

Because GLI2-ZF binding induces HDX changes at sites outside the neck-coil domain, including the microtubule-binding interface of KIF7-DM **(Figure 1)**, we used neck-coil perturbations to test whether conformational changes originating at the neck-coil can be communicated across domains to the motor. To address this, we performed HDX-MS experiments with two previously biochemically characterized mutant KIF7-DM proteins with altered neck-coil: a KIF7M-KIF27NC_chimera_, in which the KIF7DM_WT_ neck-coil is replaced with that of a closely related homolog, KIF27, and a point mutant KIF7-DM_E502A_ **(Figure 2a)**. Neither protein shows significant GLI2-ZF binding to the neck-coil, and both have a higher microtubule-binding affinity than KIF7-DM_WT_ in the absence of GLI2-ZF^3^. We compared the HDX-MS profiles of these proteins with KIF7-DM to determine whether targeted perturbations in the neck-coil domain alter motor-domain conformational dynamics.

**Figure 2.**
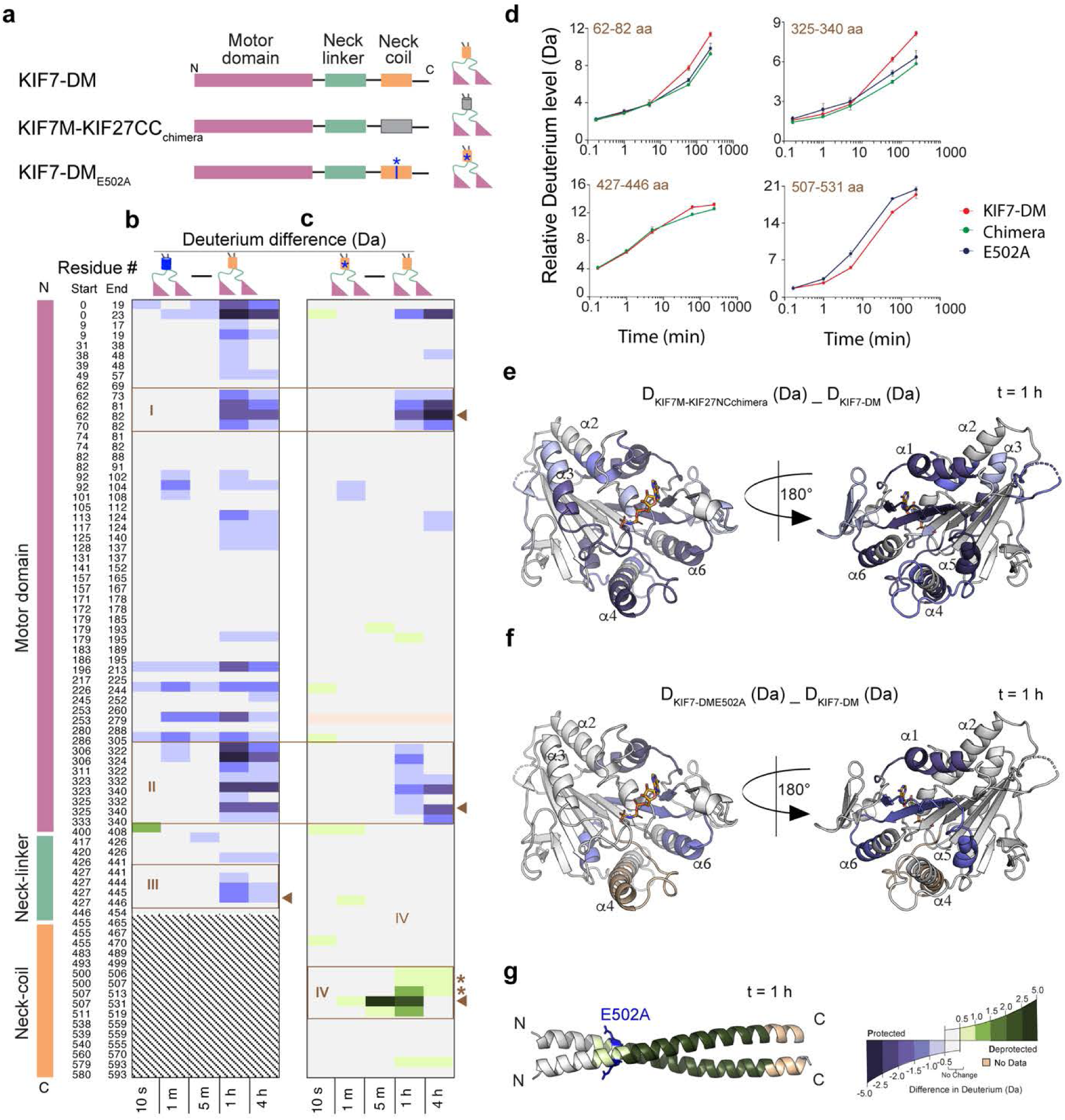
**Conformational dynamics at the KIF7 motor domain are linked to changes in its neck-coil domain** (a) Domain organization of KIF7-DM, KIF7M-KIF27NC_chimera,_ and KIF7-DM_E502A_. (b) HDX-MS data for curated peptides in KIF7M-KIF27NC_chimera_. Chiclet plot for deuterium difference (Da) in KIF7M-KIF27NC_chimera_ relative to KIF7-DM_WT_ is presented. The X-axis shows the time points, and the Y-axis shows each peptide. Only the peptides that are coincident in both proteins (KIF7-DM and KIF7M-KIF27NC_chimera_) are shown. The deuterium difference data are calculated by subtracting the absolute deuterium incorporation of the KIF7-DM from KIF7M-KIF27NC_chimera_. The differences are colored according to the scale shown. (c) HDX-MS data for curated peptides in KIF7-DM_E502A_. Chiclet plot for deuterium difference (Da) in KIF7-DM_E502A_ relative to KIF7-DM is presented. The X-axis shows the time points, and the Y-axis shows each peptide. Only the peptides that are coincident in both proteins (KIF7-DM and KIF7-DM_E502A_) are shown. The deuterium difference data are calculated by subtracting the absolute deuterium incorporation of the KIF7-DM from KIF7-DM_E502A_. The differences are colored according to the scale shown. * indicates peptides in KIF7-DM_E502A_ that contain the Ala compared to the wild-type peptide that contains Glu. (d) Deuterium incorporation curves of selected peptides indicated with solid brown arrowheads in (b) are shown. (e) Deuterium difference (Da) of the selected peptides at the 1h time point in KIF7M-KIF27NC_chimera_ in (b) are mapped on the structure of the KIF7 motor domain (PDB ID: <u>7RX0</u>) and colored according to the deuterium difference color scale. (f) Deuterium difference (Da) of the selected peptides at the 1h time point in KIF7-DM_E502A_ in (c) are mapped on the structure of the KIF7 motor domain (PDB ID: <u>7RX0</u>) and colored according to the deuterium difference color scale. (g) Deuterium difference (Da) of the selected peptides from the 1 h time point in KIF7-DM_E502A_ in (c) are mapped on the structural model of the KIF7 neck-coil and colored according to the deuterium difference color scale.

The difference HDX plot of KIF7M-KIF27NC_chimera_ relative to KIF7-DM showed extensive alterations in deuterium incorporation in both the motor domain and the neck-linker. Since the amino acid sequence of the neck-coil domain of the chimera is from KIF27 (20% identity with KIF7-NC), it was excluded from our differential MS analysis. First, we observed a region of decreased HDX in the neck-linker of the chimera (427-446 aa) **(Figure 2b III, 2e)**. Remarkably, these are the same neck-linker peptides that show decreased HDX in KIF7-DM when it is in complex with GLI2-ZF. Second, we observed decreased HDX in multiple regions of the motor domain in the chimera. These changes were: 1) in the N-terminal region of the motor domain (α1-helix 62-82 aa, P loop 92-104 aa & α2-helix 113-124 aa) **(Figure 2b I, 2e),** 2) in the major microtubule binding interface (α5-helix & α6-helix 306-340 aa) of the motor domain **(Figure 2b II, 2e)**, 3) in the switch I (196-213 aa), 4) switch II (253-279 aa), and 5) in the K-loop (286-305 aa). These data suggest that replacing the KIF7 neck-coil with the corresponding domain from KIF27 alters the conformational dynamics of the neck-linker and the microtubule-binding interface.

We next examined the point mutant KIF7-DM_E502A._ The difference HDX plot of the KIF7-DM_E502A_ relative to KIF7-DM showed increased HDX in the KIF7-DM_E502A_ neck-coil near the mutated residue **(Figure 2c IV, 2g)**. We did not observe any protection in the neck-linker of the KIF7-DM_E502A_ compared to the wild-type construct. This distinction is further evident when we compare the KIF7M-KIF27NC_chimera_ with the KIF7-DM_E502A_ **(Supplemental Data File Tab FigS2a)**. Importantly, we observed decreased HDX at the N-terminus (62-82aa) and the microtubule-binding interface of the motor domain (306-340aa) **(Figure 2c I & II, 2f**), which was also observed with the KIF7M-KIF27NC_chimera._ Thus, HDX analyses of the chimera and the point mutant collectively indicate that perturbations to the KIF7 neck-coil domain alter conformational dynamics in the neck-linker and the microtubule-binding interface of the motor domain even in the absence of GLI2.

Because GLI2-ZF binds the motor domain of KIF7-DM in addition to the neck-coil, we used the KIF7M-KIF27NC_chimera_, which lacks the neck-coil but retains the motor domain, to dissect which HDX changes in KIF7-DM upon GLI2-ZF binding arise from interactions with the neck-coil or the motor domain. For this, we compared the difference HDX-MS profiles of KIF7M-KIF27NC_chimera_ in the absence and presence of excess GLI2-ZF **(Extended Data Figure 2)**. As expected, no changes in HDX were detected at the neck-coil domain of the chimera upon GLI2-ZF binding. However, we observed increased HDX in the chimera upon interaction with GLI2-ZF at one region of the microtubule binding interface (303-322 aa) consisting of α5-helix, Loop L13, and β8 strand of the KIF7 motor domain. Strikingly, this region displayed an identical response in KIF7-DM upon GLI2-ZF binding. Hence, this localized increase in conformational dynamics can be attributed to GLI2-ZF binding to the motor domain. Notably, we did not observe any changes in HDX in the ATP-binding pocket, α4-helix and α6-helix of the microtubule-binding interface, or the neck-linker of kinesin, suggesting that these alterations to backbone dynamics in KIF7-DM arise from GLI2 binding to the neck-coil domain.

Taken together, these findings reveal that perturbations in the neck-coil domain of KIF7 are allosterically communicated to the motor domain through a long-range intramolecular conformational change, reminiscent of autoinhibition and cargo-induced activation mechanisms proposed for some motile kinesin^20–26^.

### Structural Model for KIF7-DM Autoinhibition

To gain insights into the structural basis of KIF7-DM autoinhibition, we generated a structural model of KIF7-DM using AlphaFold3 (AF3) **(Figure 3)**. The model revealed a “compact” conformation, with the two motor domains folded back onto the neck-coil domain in all five AF3 models generated **(Figure 3a)**. The complete AF3 modeling with pLDDT and PAE scores is shown in **(Extended Data Figure 3a, 3b)**.

**Figure 3.**
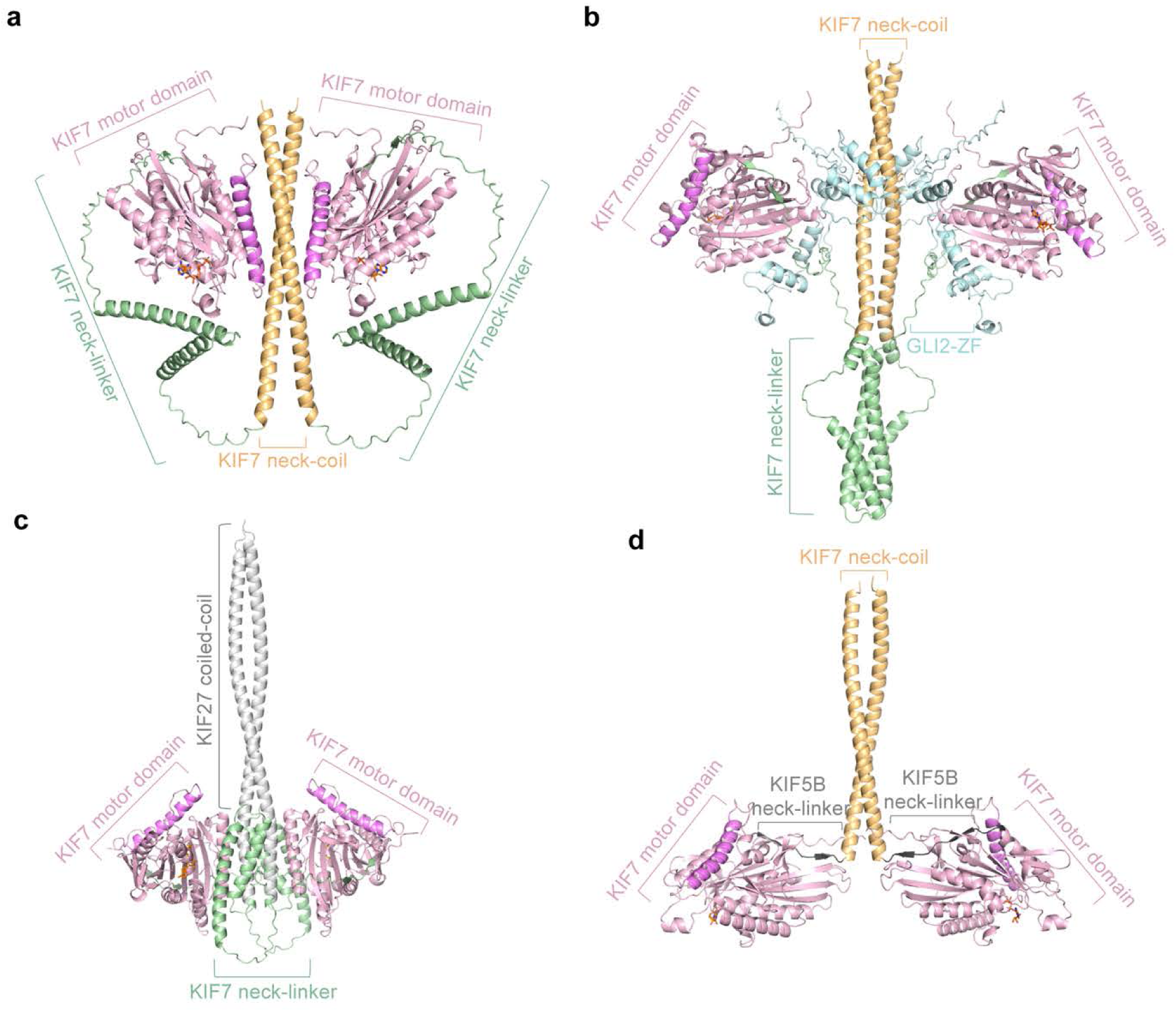
**Structural Models of Autoinhibited and Activated KIF7-DM** In all models, the motor domain is in light pink, the neck-linker in light green, the neck-coil in light orange, the swapped domains in grey, and GLI2-ZF in light blue. The α4-helix of the motor domain is colored in bright pink to represent the microtubule-binding interface of the KIF7 motor domain. (a) AF3 model of KIF7-DM (b) AF3 model of KIF7-DM with two copies of GLI2-ZF (c) AF3 model of KIF7M-KIF27NC_chimera_ (d) AF3 model of KIF7M-KIF5BNL-KIF7NC_chimera_

A striking feature of the model is that the major microtubule-interacting interface of KIF7 is in proximal to the neck-coil, which has a highly negatively charged (pI ∼4.4) surface^3^. The model suggests that this negatively charged KIF7-NC interacts intramolecularly with the microtubule-binding interface of the motor domain, which is a positively charged interface for binding the negatively charged microtubule lattice^27^. This compact conformation is consistent with the protection of the neck-coil and the microtubule-binding interface observed in KIF7-DM in the absence of GLI2-ZF in the HDX-MS experiments **(Figure 1c)**. We examined whether the monomeric KIF7 motor domain could bind the KIF7-NC by performing binding assays with purified proteins. However, we did not observe binding between these isolated protein domains in trans, suggesting either a weak affinity interaction or a specific requirement of the KIF7-NL not captured in AF3 models. Collectively, the AF3 modeling and the HDX data indicate that autoinhibition in dimeric KIF7-DM likely arises from intramolecular occlusion of the microtubule-binding interface by the neck-coil.

Additionally, the model suggests that the unusually long neck-linker of KIF7 may facilitate the intramolecular interactions between the motor and the neck-coil domains. To explore this possibility, we modeled a chimera (KIF7M-Kif5BNL-KIF7NC_chimera_) where the KIF7-NL (352-476aa) was replaced with the 12-amino acid neck-linker of conventional motile kinesin 1 (Kif5B) **(Figure 3d)**. The full AF3 model with pLDDT and PAE scores is shown in **(Extended Data Figure 3g, 3h)**. All the KIF7M-Kif5BNL-KIF7NC_chimera_ models showed an open conformation reminiscent of conventional kinesin, consistent with the idea that the long neck-linker of KIF7 may contribute to the formation of the compact, autoinhibited state.

### Structural Model for KIF7-DM Activation

We next asked how GLI2-ZF binding remodels this autoinhibited conformation to activate KIF7-DM. To examine this, we modeled the KIF7-DM:GLI2-ZF complex using AF3 **(Figure 3b)**. The full AF3 model with pLDDT and PAE scores is shown in (**Extended Data Figure 3c, 3d)**. First, across all models, the zinc fingers bind to the primary interaction site in the neck-coil domain of KIF7, consistent with the increased protection observed in HDX-MS experiments. Second, this interaction displaces the motor domains relative to the neck-coil, so that the microtubule-binding interface is no longer oriented towards the neck-coil and instead faces away from it **(Figure 3b),** consistent with changes in the HDX profile of the motor domain. Third, the model shows that GLI2-ZFs 3 and 5 are in proximity (< 6Å) to the KIF7 motor domain α2-helix and α3-helix, respectively **(Extended Data Figure 3c)**. The orientation of the zinc fingers on the motor domain differs between the AF model of the KIF7-DM:GLI2-ZF complex in the absence of microtubules and that seen in our prior EM reconstruction of the complex bound to microtubules^3^ **(Extended Data Figure 1d)**. Finally, the model shows conformational changes in the KIF7 neck-linker upon GLI2-ZF binding, forming a helical bundle. Remarkably, the helical bundle comprises peptides (417-446aa) that form the KIF7-NL_helix_ and show increased protection in HDX in the KIF7-DM:GLI2-ZF complex relative to KIF7-DM (**Figure 1c).** Together, the model suggests that GLI2-ZF binding at the neck-coil is accompanied by a conformational change in the neck-linker and a reorientation of the motor domains that relieves autoinhibition.

Since the modeling suggests a central role for the neck-coil in autoinhibition/activation of KIF7-DM, we examined the effects of changing the KIF7-NC by AF3 modeling of the constitutively active KIF7M-KIF27NC_chimera_ **(Figure 3c)**. The full AF3 model with pLDDT and PAE scores is shown in **(Extended Data Figure 3e, 3f)**. All the models revealed an “open” conformation, distinct from KIF7-DM and reminiscent of the dimeric motile kinesin^28^. In this structure, the motor domains are displaced from the neck-coil and repositioned toward the neck-linker, leaving the microtubule-binding interface accessible for microtubule binding **(Figure 3c)**. Additionally, the helices in the KIF7-NL fold into a compact bundle close to the N-terminal region of the KIF7-NC (427-446aa) **(Figure 2b),** the same region that forms a helical bundle in the KIF7-DM:GLI2-ZF complex. Notably, the KIF7-NL_helix_ and the motor domain are also regions where the HDX profile of the chimera differs significantly from KIF7-DM. Thus, reorientation of the motor domain away from the neck-coil and remodeling of the neck-linker helix emerge as shared features of the activated state of KIF7-DM.

Collectively, the modeling provides a structural framework that contextualizes the HDX-MS findings in 3-D space. It positions the neck-coil as a regulatory element that can occlude the microtubule-binding interface of the KIF7 motor domain. Either GLI2-ZF binding or modifications to the neck-coil sequence displace the motor domains, rendering the microtubule-binding interface more accessible. In both cases, this transition is accompanied by a conformational change in the KIF7 neck-linker.

### Insights into KIF7 autoinhibition from comparative analyses of the neck-coil domain

The AF3 modeling suggests that a closed conformation, in which the motor domain is positioned close to the neck-coil of KIF7, underlies its autoinhibition. This model makes two predictions. First, it predicts that in the absence of the motor domain, the isolated neck-coil would be less protected and exhibit higher HDX than when the motor domain is present. Second, it predicts that changes in the neck-linker upon GLI2 binding will occur at different kinetics for an isolated neck-coil versus the KIF7-DM, due to competing interactions with the motor domain.

To test the first prediction, we performed HDX-MS on a previously characterized recombinant isolated neck-coil domain dimer (KIF7-NC_isolated_) and compared it with the neck-coil domain in the intact KIF7-DM (KIF7-NC-DM) **(Figure 4)**. Since the neck-coil domains from these two proteins did not yield the same digestion profiles, coincident peptides cannot be obtained for direct comparisons. Therefore, we have presented these data as relative percent deuterium incorporation for all peptides in a linear fashion for the KIF7-NC region (aa 490-550) **(Figure 4).** Comparison of the HDX profiles between KIF7-NC_isolated_ and KIF7-NC-DM revealed two major differences. First, proteolytic digestion of KIF7-NC_isolated_ yielded 100% sequence coverage with 8.40 redundancy, while that for KIF7-NC-DM yielded 88.3% coverage with only 2.56 redundancy **(Supplementary Data File Tab 2).** This suggests distinct acid-induced denatured states between the isolated neck-coil domain than that in the larger construct. Second, the isolated KIF7-NC domain exhibited rapid deuterium incorporation, reaching more than 60% relative deuterium uptake within 5 minutes, suggesting a highly dynamic and unstable secondary and/or tertiary structure (**Figure 4a-b)**. In contrast, the neck-coil within the KIF7-DM showed markedly lower deuterium incorporation, ranging between 10–40% at 5 minutes and approaching ∼60% relative deuterium incorporation after 1 hour of exposure **(Figure 4c-d**). Together, these data indicate that the KIF7 neck-coil is markedly stabilized in the context of the intact KIF7-DM, consistent with restricted conformational dynamics in the autoinhibited state.

**Figure 4.**
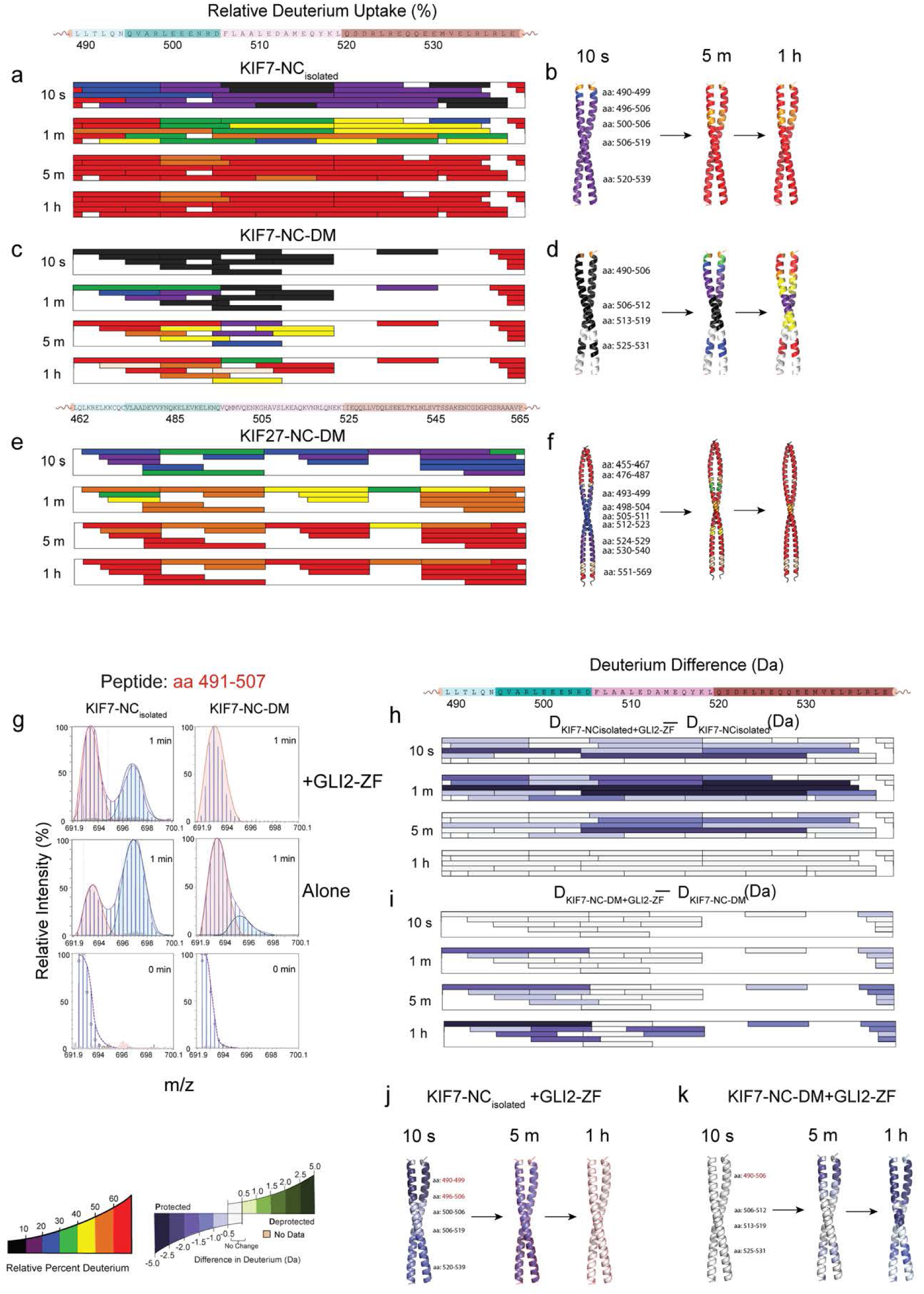
**Insights into KIF7 autoinhibition from comparative analysis of HDX-MS of the neck-coil domain in different contexts** (a) The linear map of KIF7-NC (489-539 aa residues) was divided into 4 peptide segments for comparison and aligned to the chiclet plot of the relative percentage of deuterium uptake for KIF7-NC_isolated_ peptides. The level of relative deuterium incorporation %D for each time point is rendered in colored horizontal layers for each peptide, allowing for visualization of the rate of exchange in each segment. The gaps represent the amino acid sequence in which no information was obtained due to incomplete coverage of identical peptides. (b) Chiclet plot of the relative percentage of deuterium uptake for KIF7-NC-DM peptides. (c) The linear map of the KIF27-NC (462-565 aa residues) was divided into 4 peptide segments for comparison and aligned to the chiclet plot of the relative percentage of deuterium uptake for KIF27-NC-DM peptides. (d) The relative deuterium incorporation (%D) for KIF7-NC_isolated_ at 10 s, 5 m, and 1 h, mapped onto the homology-modeling derived structure of KIF7-NC. (e) The relative deuterium incorporation %D for KIF7-NC-DM at 10 s, 5 m, and 1 h mapped onto the homology-modeling derived structure of KIF7-NC. (f) The relative deuterium incorporation %D for KIF27-NC-DM at 10 s, 5 m, and 1 h mapped onto the homology-modeling derived structure of KIF27-NC. (g) Representative spectra for neck-coils, KIF7-NC_isolated_ and KIF7-NC-DM, showing mass shifts for peptide 491-507, which shows a bimodal distribution, with increasing deuteration time. The two populations are colored red and blue. (h) The differential deuterium (Da) for KIF7-NC_isolated_ in the presence and absence of GLI2-ZF. Data for each peptide is shown at different time points. and is rendered in colored horizontal layers according to the legend below the figure. (i) The differential deuterium (Da) for KIF7-NC-DM in the presence and absence of GLI2-ZF is displayed for each peptide at each time point and is rendered in colored horizontal layers according to the legend below the figure. (j) The deuterium difference (Da) from (h) is mapped on the homology modelling structure of KIF7-NC at three time points (10 s, 5 m, and 1 h). (k) The deuterium difference (Da) from (i) is mapped on the homology modelling structure of KIF7-NC-DM at three time points (10 s, 5 m, and 1 h).

To distinguish motor-dependent stabilization from intrinsically lower coiled-coil stability of the isolated neck-coil, we compared HDX of the KIF27 neck-coil in the KIF7M-KIF27NC_chimera_ (KIF27-NC-DM), which provides a similarly sized coiled-coil predicted not to interact with the motor domain, with KIF7-NC_isolated_ and the KIF7-NC-DM. If the high HDX observed for KIF7-NC_isolated_ reflected generic instability of an isolated coiled-coil, we would expect the KIF27 neck-coil in the chimera to show lower HDX levels similar to the neck-coil in KIF7-DM. For the comparison of the HDX profiles between the KIF7-NC_isolated_ and KIF27 neck-coil in the chimera, we examined the peptides of the neck-coil spanning amino acids 462–565 for KIF27 since the protein sequence is different from KIF7 **(Figure 4e-f)**. These results show that KIF27-NC-DM exhibited rapid deuterium incorporation, with more than 20% labeling at 10 s, and 40–50% labeling by 1 min **(Figure 4e)**. Similar to the KIF7-NC_isolated_, deuterium incorporation was greater than 60% within 5 mins **(Figure 4f)**. These findings suggest that, despite being a dimeric kinesin context, the neck-coil of KIF27 is highly dynamic, similar to KIF7-NC_isolated_. Therefore, the protection of the neck-coil in KIF7-DM is likely to arise from its structural context rather than intrinsic coiled-coil stability.

To test the second prediction and interrogate the compact conformation in an independent experiment, we examined how GLI2-ZF binding alters neck-coil dynamics in the absence versus presence of the motor domain. Therefore, we generated two SEC-purified complexes, KIF7-NC_isolated_:GLI2-ZF and KIF7-DM:GLI2-ZF, and examined HDX-MS of these and the corresponding KIF7 proteins alone **(Figure 4h-k)**. We found that the raw MS data for KIF7-NC_isolated_ showed non-EX2 EX1-like bimodal distributions^18^ for most peptides (aa 490-499, 490-505, 490-506, 500-507, 507-524, 525-531, 532-539), both in the absence and presence of GLI2-ZF **(Supplementary Data File Tab EX1_KIF7 NC).** As shown in **Figure 4g**, the raw m/z data shows two distinct isotope clusters for the same peptide, with the lower-mass envelope (red, population 1) representing an exchange-incompetent population and a higher-mass envelope (blue, population 2) corresponding to a more exchange-competent state. We observed that GLI2-ZF binding reduced the likelihood of the more open exchange-competent state in KIF7-NC_isolated_ (population 1; from 60% in KIF7-NC_isolated_ to 100% in KIF7-NC_isolated_:GLI2-ZF) but did not fully restore it to the protected state. These findings are consistent with stabilization of the neck-coil by GLI2-ZF binding (**Figure 4g, Supplemental data file tab EX1)**.

Differential HDX of KIF7-NC_isolated_:GLI2-ZF and KIF7-NC_isolated_ shows that, when bound to GLI2-ZF, the neck-coil exhibited low HDX at the earliest labeling time point of 10 s, which was further reduced at 1 min. However, this protection progressively diminished by 5 min and was absent at 1 h **(Figure 4h, 4j).** To examine how the presence of the motor domain alters this HDX profile, we compared it to the differential HDX of KIF7-DM:GLI2-ZF and KIF7-DM. We observed that the neck-coil in the context of the KIF7-DM:GLI2-ZF also showed decreased HDX when compared to KIF7-DM **(Figure 4i, 4k)**. However, unlike in the isolated neck-linker, no change in HDX was detectable at the earliest labeling time point of 10 s. Reduction in HDX was evident by 1 min of deuterium labeling, and became more pronounced by 5 min, and was further enhanced by 1 h **(Figure 4k)**. These findings reveal a time lag and slower overall kinetics in the protection of the neck-coil by GLI2-ZF binding in the context of KIF7-DM, compared to KIF7-NC_isolated_, suggesting that the environment of the neck-coil is altered in the presence of the neck-linker and the motor domains.

Collectively, these two sets of experiments suggest that in KIF7-DM, the neck-coil is in a more protected environment compared to the isolated neck-coil or in the chimera with the KIF27 neck-coil. These findings are consistent with the AF3 modeling, where the compact state is observed only in the context of the KIF7-DM.

### ATP Turnover Regulates the Function of the Non-motile KIF7

In addition to autoinhibition, kinesin–microtubule interactions are strongly influenced by nucleotide state, raising the question of how the low but measurable ATPase activity of the non-motile kinesin KIF7 contributes to its microtubule binding-unbinding dynamics.

Findings from the HDX data **(Figure 1c-g)** indicate that GLI2 binding affects the structural dynamics of the KIF7 ATP pocket and increases the KIF7 ATPase activity **(Extended Data Figure 1c)**. However, since KIF7 is non-motile and has significantly reduced microtubule-stimulated ATPase activity compared to conventional kinesins, the importance of KIF7’s ATPase activity for its function is unclear. To investigate this, we generated an ATPase-deficient “rigor” mutant by mutating a conserved Lys residue in the Walker A motif (P-loop) to Ala (K100A) **(Figure 5a)**. This KIF7-DM_K100A_ mutant protein is well-folded, with a size-exclusion chromatography profile similar to that of the KIF7-DM protein **(Extended Data Figure 4a,b)**. KIF7-DM_K100A_ had no detectable microtubule-stimulated ATPase activity compared to KIF7-DM **(Figure 5b)**. Next, we compared the microtubule-binding of wild-type and K100A mutant proteins using an *in vitro* total internal reflection fluorescence microscopy assay^3^ **(Extended Data Figure 4c,d)**. At a fixed KIF7 concentration of 100 nM, we found a >10-fold increase in the intensity per unit length of KIF7-DM_K100A_ relative to KIF7-DM **(Figure 5c)**. These data validate that KIF7-DM_K100A_ has lower ATPase and higher microtubule high-affinity compared to the wild-type protein.

**Figure 5.**
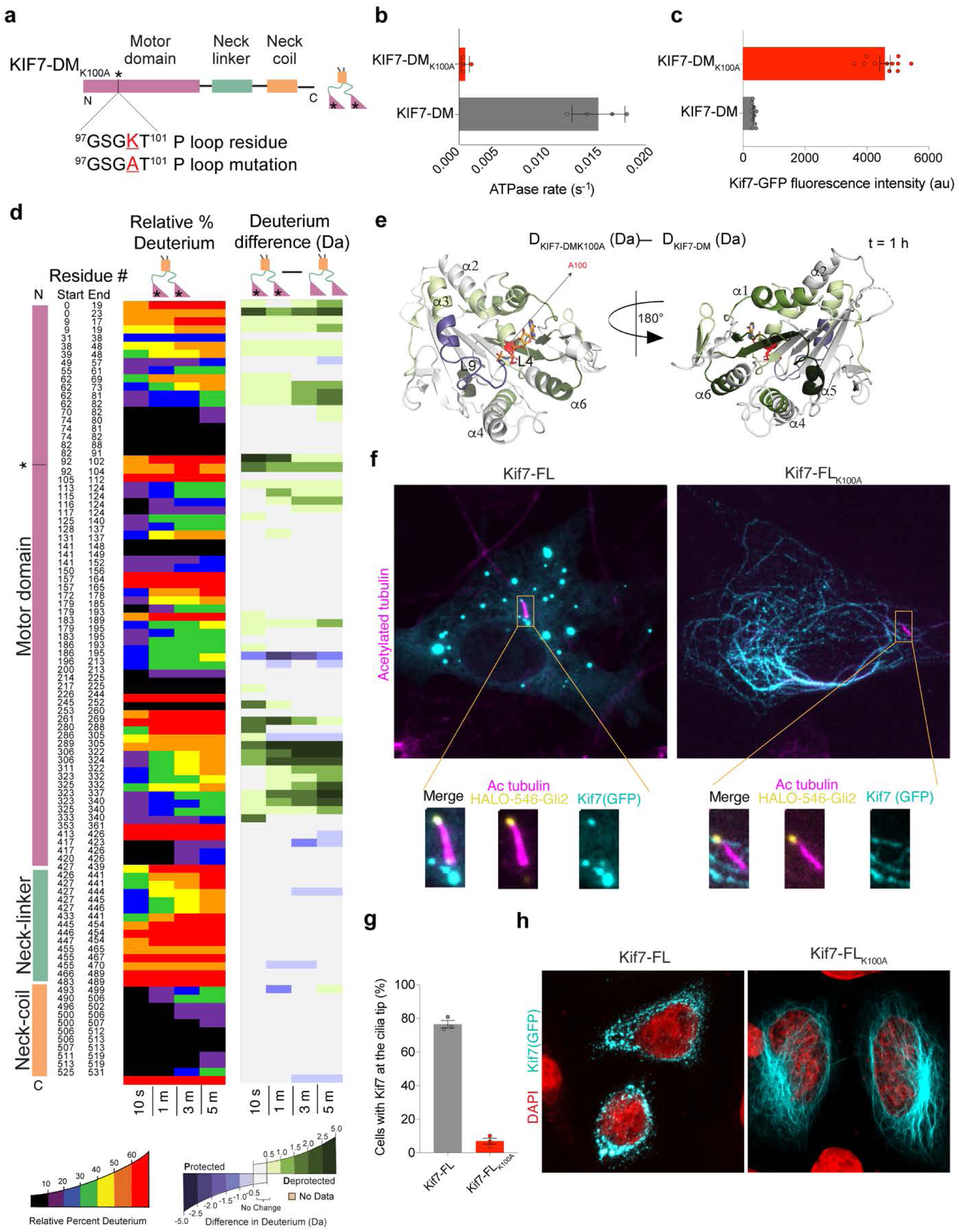
**ATP turnover regulates the function of the non-motile KIF7** (a) Domain organization of KIF7-DM_K100A_ showing the residue mutated. (b) Microtubule-stimulated ATPase of 1 µM KIF7-DM_K100A_ and 1 µM KIF7-DM (n=3). (c) Analysis of KIF7-GFP fluorescence intensity per unit microtubule length from TIRFM assays with 100 nM KIF7-DM_K100A_ and 100 nM KIF7-DM. (d) HDX-MS data for curated peptides in KIF7-DM_K100A_. Chiclet plots for relative percentage deuterium for KIF7-DM_K100A_ (left column) and deuterium difference between KIF7-DM_K100A_ and KIF7-DM (right column) are presented. The X-axis shows the time points, and the Y-axis shows each peptide. Only the coincident peptides in KIF7-DM_K100A_ and KIF7-DM are shown. The deuterium difference data are calculated by subtracting the relative deuterium incorporation of the KIF7-DM from the KIF7-DM_K100A_. The differences are colored according to the scale shown. “*” indicates peptides in KIF7-DM_K100A_ that contain Alanine instead of Lysine. (e) Deuterium difference of the selected peptides from 1 h time point in KIF7-DM_K100A_ is mapped on the structure of the KIF7 motor domain (PDB ID: <u>7RX0</u>) with the deuterium difference color scale shown. (f) Immunofluorescent images of HALO-Gli2 expressing NIH3T3 cells transfected with GFP-tagged Kif7-FL or Kif7-FL_K100A_ constructs (cyan). The ciliary axoneme is visualized using Acetylated tubulin (magenta), and HALO-Gli2 is visualized using the HALO-546 dye (yellow). Images were acquired 12 h after pathway activation using the HH pathway agonist (SAG). (g) Percentage of cells from experiments as in (f) that show Kif7-GFP signal at the cilium tip 24 hours after SAG treatment. (h) HeLa cells expressing GFP-tagged Kif7-FL or Kif7-FL_K100A_ constructs (cyan). DAPI was used to stain the nucleus (red).

We next performed HDX-MS analysis of the KIF7DM_K100A_ mutant to examine how this single point mutation in the ATPase pocket affects its structural dynamics. The digestion profile of the KIF7-DM_K100A_ mutant showed a higher sequence coverage (>90%) compared with KIF7-DM (82.9%) **(Supplemental data file tab 2)**. This higher digestibility suggests that the KIF7-DM_K100A_ acid-denatured state is conformationally different from wild type. Comparative HDX analysis of KIF7-DM_K100A_ to KIF7-DM revealed no significant changes in HDX within the neck-coil and neck-linker regions between the two proteins. In contrast, we unexpectedly observed extensive increases in HDX across multiple areas of the motor domain relative to wild type. This increase in HDX was observed from the earliest time point examined (10 s), suggesting increased conformational flexibility. The specific regions with observed increased deuterium incorporation included the ATP binding pocket in the P-loop (92-104aa), and the microtubule binding interface (280-332aa), which is comprised of helices α4, α5, α6, and the K-loop **(Figure 5d , 5e)**. The only region in the motor domain that showed decreased levels of deuterium incorporation, compared to KIF7-DM, was in switch I (196-213aa) **(Figure 5d , 5e)**.

We used the K100A mutant to understand the relevance of KIF7 ATPase activity to its cellular function. We transiently overexpressed GFP-tagged full-length wild-type mouse Kif7 (Kif7-FL) and K100A (Kif7-FL_K100A_) in NIH3T3 cells stably expressing HALO-Gli2. We examined how reduced Kif7 ATPase activity and increased microtubule binding of the mutant affect Kif7 localization to the distal cilium tip in response to Hh activation by the pathway agonist SAG **(Figure 5f)**. Acetylated tubulin antibodies marked the cilium shaft, and HALO-546 dye marked Gli2 at the cilium tip. The cilium tip localization of Gli2 was not compromised, as these cells express endogenous wild-type KIF7. We found that <10% of Kif7-FL_K100A_ expressing cells have a GFP signal at the cilium tip, as compared to Kif7-FL, which accumulated at cilia tips upon Hedgehog activation in ∼78% of cells **(Figure 5g)**. In the Kif7-FL_K100A_ cells, we observed a striking localization of the mutant Kif7 protein on cytoplasmic microtubules, which was not seen for the autoinhibited wild-type Kif7 (as noted in the introduction, full wild-type Kif7 is more autoinhibited than KIF7-DM). HALO-546-Gli2 was also enriched on these microtubules in the K100A mutant. These data indicate that the mutant kinesin is sequestered on cytoplasmic microtubules, which hinders its trafficking to the cilium tip. We also validated the cytoplasmic microtubule sequestration of transiently expressed Kif7-FL_K100A_ in HeLa cells **(Figure 5h).** These observations indicate that loss of ATP turnover traps Kif7 on cytoplasmic microtubules, preventing efficient trafficking to the cilium tip.

In summary, the ATPase-deficient K100A mutant of KIF7 exhibits increased microtubule affinity and motor-domain conformational dynamics *in vitro*, and defects in ciliary localization in cells. Thus, ATP hydrolysis, even in a non-motile kinesin, plays a role in dynamic release from microtubule to avoid cytoplasmic sequestration and enable the localization of KIF7 to the cilium tip upon pathway activation.

## Discussion

Our study reveals how canonical regulatory principles of motile kinesins—autoinhibition, activation through cargo binding, and ATPase-linked microtubule interactions—have been repurposed in KIF7 for its motility-independent role as a cilium-tip scaffold during Hedgehog signaling.

Based on combined HDX-MS and AF analyses, we propose a mechanism for KIF7 autoinhibition and its reversal by GLI2 binding **(Figure 6)**. In the autoinhibited state, the KIF7 neck-coil occludes the microtubule-binding interface of the motor domain, stabilizing a compact conformation with restricted microtubule access. The unusually long neck-linker of KIF7 enables this intramolecular interaction by allowing the N-terminal motor domain to fold back onto the C-terminal neck-coil. GLI2 binding to the neck-coil disrupts this interaction, leading to coordinated conformational changes across the neck-coil and neck-linker and extending into the motor domain, including the microtubule-binding interface and ATPase pocket, thereby relieving autoinhibition. Thus, KIF7 preserves a core kinesin regulatory logic—autoinhibition mediated by intramolecular interactions between microtubule-binding and cargo-binding elements—while implementing it through a distinct structural solution. The highly acidic neck-coil, previously shown to engage the positively charged GLI2 zinc-finger domain via DNA-mimicry principles, is repurposed here as an autoinhibitory docking site for the positively charged microtubule-binding surface of the motor domain. In this way, electrostatic complementarity, which promotes kinesin–microtubule interactions, is co-opted in KIF7 to stabilize an autoinhibited state.

**Figure 6.**
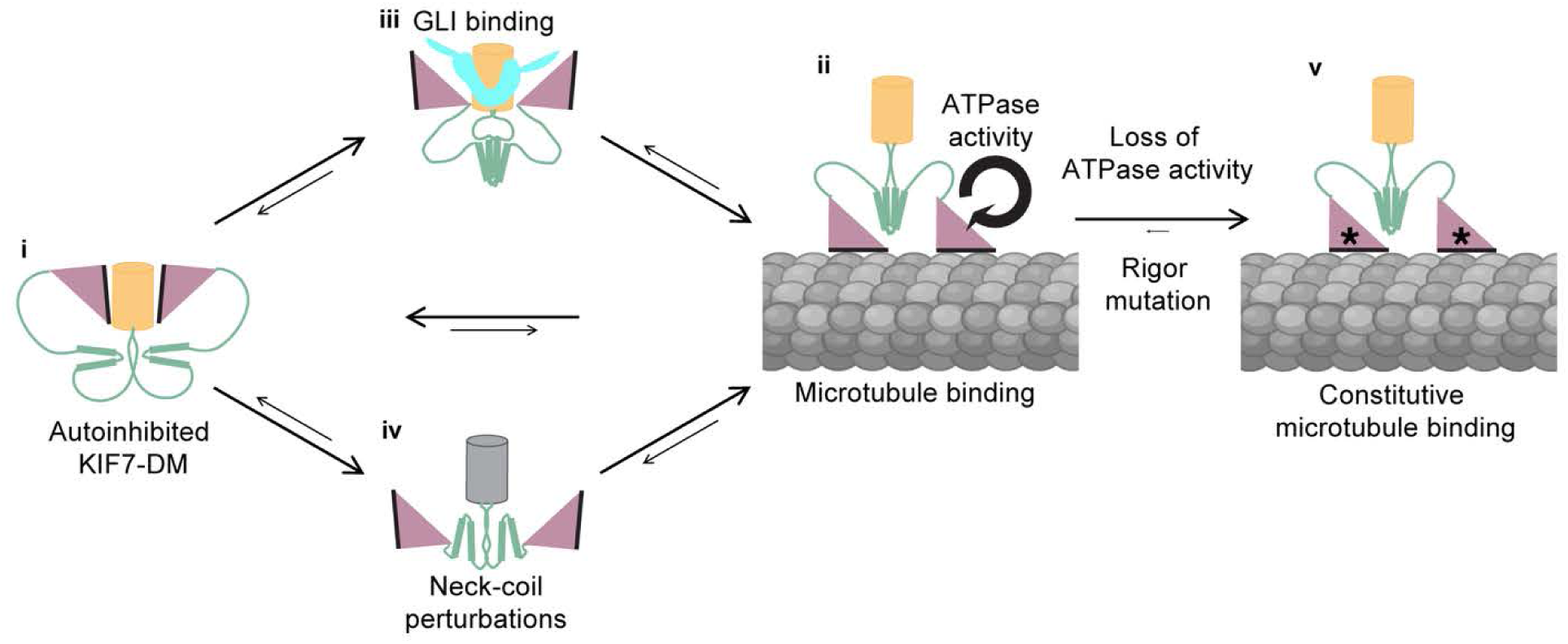
**Mechanism of KIF7-DM autoinhibition and activation** Schematic representation of the proposed mechanism. i. In the autoinhibited state, the microtubule-binding interface (black) of the KIF7-DM motor domains (pink) is held proximal to the neck-coil domain (orange). The long and flexible neck-linker (green) promotes the intramolecular interaction between motor and neck-coil domains. ii. The autoinhibited state is in dynamic equilibrium with the active microtubule-bound state. iii. GLI2-ZF (blue) binding promotes the activated state by displacing the KIF7 motor domains from the neck-coil such that the microtubule-binding interface faces away from the neck-coil, increasing the probability of interacting with microtubules. Additionally, the neck-linker helix reorganizes into a helical bundle. iv. Perturbations to the neck-coil similarly displace the motor domain to promote an open microtubule binding compatible conformation. v. An ATPase-deficient rigor mutation shifts the equilibrium, resulting in constitutive microtubule binding. Thus, the low but measurable ATPase activity of this non-motile kinesin ensures that KIF7-DM is recycled and remains in equilibrium with the autoinhibited state.

Independent HDX-MS measurements support the proposed autoinhibition and activation mechanism and, together with AF models, define four features underlying KIF7 regulation. First, in the absence of GLI2-ZF, the neck-coil in KIF7-DM displays a significantly lower level of HDX compared to either the isolated neck-coil or the neck-coil in the KIF7M–KIF27NC chimera, indicating that KIF7’s motor domain shields the neck-coil in the autoinhibited state. Second, GLI2-ZF binding further lowers the level of HDX of the neck-coil in KIF7-DM, but with slower kinetics than observed for GLI2 binding to the isolated neck-coil. This delay is consistent with motor-domain displacement from the neck-coil preceding GLI2-ZF binding in the autoinhibited KIF7-DM. Third, activation is accompanied by conformational changes within the neck-linker helix, which exhibits significant decreases in HDX in two different activated states (KIF7-DM:GLI2-ZF and KIF7M–KIF27NC chimera), identifying it as a structural element that undergoes rearrangement upon relief of autoinhibition. Finally, HDX-MS reveals changes in the conformational dynamics of the motor microtubule-binding interface in both activated states, relative to KIF7-DM alone, which correlate with greater microtubule-binding affinity. AF models provide a predictive structural framework for these observations, visualizing a compact, autoinhibited conformation of KIF7-DM in which the motor microtubule-binding surface is positioned against the acidic neck-coil, as well as open, activated states in which this interaction is disrupted, and accompanied by reorganization of the neck-linker helix. Together, these observations provide insights into how GLI2 binding induces long-range allosteric conformational changes in KIF7, shifting it from a compact, autoinhibited state to an open, microtubule-binding configuration.

Kinesins share a modular autoinhibition mechanism but execute it using family-specific architectures. In Kinesin-1, the C-terminal tail folds over to make intramolecular contacts with the motor domains, which are further stabilized by the light chains, locking it in an inactive, non-motile state^20,21,24,29^. In Kinesin-2, the motor domains are folded back and sequestered against the distal stalk and stabilized by a β-hairpin^26^ and/or phosphorylation^22,23^, occluding the microtubule-binding surface. In Kinesin-3, distal coiled coils fold onto the motor-neck assembly to form a self-inhibited monomer that is activated upon dimerization^25^. KIF7 adapts this same core principle—autoinhibition via a cargo-binding domain—but with a unique structural solution. Its compact autoinhibited state is enabled by two key evolutionary adaptations. First, the KIF7-NC domain is highly negatively charged, with a pI of ∼4.4, providing charge complementarity for binding the positively charged microtubule-binding surface of the motor domain. Second, an unusually long (>120aa) and mostly disordered neck-linker connects the KIF7-motor and the KIF7-NC. This contrasts sharply with the relatively small (∼15 amino acids) neck-linker in most kinesins, where the short length is important for the strict coordination of motor heads for processive stepping^12^. Indeed, replacing the KIF7-NL with a short 12 amino acids long neck-linker of Kif5b in the AF modeling results in an ‘open’ conformation. This unique architecture in KIF7 results in an activation mechanism where GLI relieves inhibition by engaging this same acidic neck-coil, as well as a second site on the motor domain, rather than a distal C-terminal cargo binding site. Together with defective mechanochemistry^10,11^, this retools the kinesin architecture, such that it is optimized for regulated, signal-dependent microtubule binding instead of motility.

Since the nucleotide state of KIF7 has small but significant effects on microtubule binding affinity^11^, we examined whether it contributes to KIF7’s microtubule binding-unbinding equilibrium in addition to the regulation by GLI. We characterized KIF7-DM_K100A_, a rigor mutant, which bound microtubules with higher affinity than KIF7-DM. HDX-MS of this single-point mutant reveals conformational dynamics distinct from those of the other active states we examined here. Cellular studies revealed that the rigor mutant of KIF7 remains bound on cytoplasmic microtubules and is not properly localized to the cilium after pathway activation. Therefore, ATP hydrolysis enables dynamic cycling of KIF7 on and off microtubules, facilitating its cilium tip accumulation during Hh pathway activation (**Figure 6a iv-v)**. The requirement for ATPase activity, despite KIF7 being non-motile, points to a gating mechanism in which nucleotide cycling during the ATP hydrolysis cycle allows regulatable microtubule binding.

Despite retaining a conserved kinesin molecular architecture, KIF7 functions more as a microtubule-associated protein (MAP) than an active motor. Remarkably, many aspects of canonical kinesin activity and regulation are repurposed for KIF7 function as a MAP. First, reminiscent of kinesins, autoinhibition occurs through an interaction between motor and cargo binding domains, which allows for activation through cargo binding. In contrast, MAP-microtubule interactions are mainly regulated by post-translational modifications (PTMs) or intermolecular protein-protein interactions^30^. One exception is the MAP EB1, which has been proposed to be autoinhibited by occlusion of the CH2 domains that bind microtubules^31–33^. Second, the ATPase activity is used for cycling the protein on and off microtubules. This allows a dynamic equilibrium between autoinhibited and uninhibited states, which can be exploited for signal-dependent regulation by GLI factors. This mode of regulating MT binding is not available in MAPs. Third, the conformational changes in the kinesin neck-linker are tightly coupled to conformational changes in motor domains for stepping. In KIF7, neck-linker conformational changes are repurposed for communication between the motor and the neck-coil domains. Through this re-engineering of conserved kinesin features, KIF7 functions as an HH-responsive structural scaffold at the cilium tip. It will be interesting in the future to see if the conserved kinesin features are similarly repurposed for non-motile functions, such as cell signaling for Kif26A/B^34,35^ or organizing a motile cilia scaffold in the case of KIF27^36^.

## Acknowledgements

R.S. was supported through the American Cancer Society (Ellison Foundation Research Scholar) and NIH (NIGMS) 1R01GM145651. F.H. was supported through Massachusetts General Hospital’s Executive Committee on Research Fund for Medical Discovery (ECOR-FMD) fellowship in Fundamental Research.

## Data Availability

All HDX-MS data have been deposited to the ProteomeXchange Consortium via the PRIDE^37^ partner repository with the dataset identifier **PXD074139.**

## Methods

### Experimental Model and Subject Details

Recombinant proteins were overexpressed in Sf9 cells in accordance with the Bac-to-Bac Baculovirus expression system. Recombinant proteins expressed in E. coli Rosetta (DE3) cells were induced with the addition of 1 mM IPTG after an OD of 0.6. Cells were grown for 18 hours at 4°C. Transient protein expression in NIH3T3 and HeLa cells has been detailed in the section headed “Cell Culture and Immunofluorescence”.

### Protein Expression and Purification

The N-terminal fragment of human KIF7 (Uniprot Q2M1P5) (KIF7-DM 1-543aa) was cloned into a pFastBac expression vector (Thermo) that included a tobacco etch virus (TEV) protease cleavable N-terminal 6x His-tag and SUMOstar solubilization tag and a C-terminal GFP tag. To design the KIF7-KIF27 chimera, we performed homology modeling of the two kinesins using PROMALS 3D (UTSW). We complimented this with modeling KIF7 and KIF27 CC domains using COILS (EMBL) and PairCoil (MIT) for accurate neck coil substitution. The final sequence for KIF7M-KIF27-CC chimera is KIF7 1-462aa (motor) and KIF27 475-570aa (neck coil). Dimeric KIF7 N-terminal constructs: KIF7-DM (1-543aa), KIF7-DM_E502A_, KIF7M-KIF27NC chimera and KIF7-DM_K100A_ constructs were expressed in SF9 insect cell line using the Bac-to-Bac® Baculovirus Expressions System (Thermo) with cells grown in HyClone CCM3 SFM (GE Life Sciences) and expressed from P3 virus for 72 hat 27°C. For the dimeric KIF7 N-terminal constructs, cell pellets KIF7were lysed by short sonication in buffer A (20mM PIPES pH 7.4, 150mM NaCl, 10% glycerol, 1mM MgCl_2_ and 25mM imidazole) supplemented with 100μM ATP, 2mM TCEP, 1mM PMSF, 75 U benzonase and 1X HALT (Thermo). Lysate was cleared by ultracentrifugation, and supernatant was incubated with Ni-NTA for 1H. Resin was washed with buffer A supplemented with 20μM ATP and 0.5mM TCEP and eluted with 400mM imidazole with 100μM ATP. Peak fractions were pooled and, if needed, cleaved overnight by TEV (1/30 w/w) at 4°C. Proteins were further purified by size exclusion chromatography (Superdex 200 10/300GL) in 20mM PIPES pH 7.4 150mM NaCl 5% glycerol 5mM ß-Me 1mM MgCl_2_ and 100μM ATP and frozen in liquid nitrogen.

The neck-coil dimerization domain fragment of KIF7, (KIF7-NC 460-600aa) was cloned into a modified pGEX vector that contained a human rhinovirus (HRV) protease cleavable N-terminal GST-tag. KIF7-NC was expressed in BL21 (DE3) Rosetta (Millipore) E. coli at 18°C with 0.25mM IPTG for 18-20hr. The cell pellets were lysed by short sonication in buffer B (20mM PIPES pH 7.4, 10% glycerol) supplemented with 2mM TCEP, 1mM PMSF, 75 U benzonase and 1X HALT (Thermo). Lysate was cleared by ultracentrifugation, and supernatant was incubated with GST-4B beads (GE Life Sciences) for 1H. Resin was washed with buffer B supplemented with 0.5mM TCEP and eluted with 10mM reduced glutathione in 20mM PIPES pH 7.4 and 150mM NaCl. Peak fractions were pooled and, if needed, cleaved overnight by GST-HRV-3C (1/40 w/w) at 4°C. Proteins were further purified by size exclusion chromatography (Superdex 200 10/300GL) in 20mM PIPES pH 7.4, 150mM NaCl, 5% glycerol 2mM TCEP and frozen in liquid nitrogen.

The zinc-finger domain of GLI2 (Uniprot P10070) (GLI2-ZF 418-604aa) was cloned into:

1. A modified pET-Duet-1 expression vector with an HRV protease cleavable N-terminal 6x His-tag. GLI2-ZF protein was expressed in BL21 (DE3) Rosetta (Millipore) E. coli at 18°C with 0.25mM IPTG for 18-20hr. GLI2-ZF pellets were lysed by short sonication in buffer C (20mM PIPES pH 7.4 150mM NaCl) supplemented with 2mM TCEP, 1mM PMSF, 300 U benzonase and 1X HALT (Thermo). Lysate was cleared by ultracentrifugation, and supernatant was incubated with Ni-NTA for 1H. Resin was washed with buffer A supplemented with 0.5mM TCEP and eluted with 400mM imidazole. Peak fractions were pooled and, if needed, cleaved overnight by GST-HRV-3C (1/40 w/w) at 4°C. Proteins were further purified by size exclusion chromatography (Superdex 75 10/300GL and Superdex 200 10/300GL) in 20mM PIPES pH 7.4, 150mM NaCl, 10% glycerol, 2mM TCEP and frozen in liquid nitrogen.

All proteins were greater than 95% pure and eluted as a single peak from the size exclusion chromatography column at the expected volumes for their size and oligomerization state.

### Hydrogen-Deuterium Exchange Mass Spectrometry (HDX-MS)

Each experiment was performed in duplicate. Proteins used for the experiment were diluted to 40 µM stock solutions for HDX-MS analysis. Deuterium labelling was initiated by diluting 1µL of 40 µM protein solution into 18 µL (19-fold dilution) of D_2_O buffer (20 mM HEPES, 150 mM NaCl, pD 7.4, 99% D_2_O). Experiments were performed at 20 °C. At each labelling time points (10s, 30s, 1m, 3m, 5m and 1 h), the labelling reaction was quenched with the addition of an equal volume (19 µL) of ice-cold quenching buffer (5 M guanidium hydrochloride, 0.7 M TCEP, 0.8 % (v/v) formic acid pH 2.1, H_2_O). The quenched protein samples were then subjected to an online proteolytic digestion using Affipro Nepenthesin-II column (AffiPro, AP-PC-004) at 15 °C. The resulting peptides were trapped and desalted on a VanGuard Pre-column trap (Waters, 186003980) for 3 min at 100 μL/min and separated using ACQUITY UPLC HSS T3, 1.8 μm, 1.0 mm × 50 mm column (Waters, 186003535). The trap column and the analytical column were housed in a Waters HDX cooling chamber that was held at 0 °C. The peptides were analyzed immediately using Synapt G2Si mass spectrometer operated in HDMS^E^ mode. Replicates of undeuterated control samples were used for peptide identification using PLGS 3.0.3 (Waters Corporation) as described in the Supplementary Data. The peptides identified in PLGS were then analyzed in DynamX 3.1.1 (Waters Corporation), implementing a minimum products per amino acid cutoff of 0.25 and at least two consecutive product ions (see Supplementary Data). Those peptides that passed the filtering criteria were further processed, and the relative amount of each peptide was determined using DynamX 3.1.1 by subtracting the centroid mass of the undeuterated form of each peptide from the deuterated form, at each time point, for each condition. These deuterium uptake values were used to generate all uptake graphs and difference maps. The error of determining the average deuterium incorporation for each peptide was set at ± 0.5 Da. Deuterium incorporation levels were not corrected for back exchange, and thus the data are reported as relative exchange^38^.

### AlphaFold

AlphaFold models were generated for the various protein sequences using Alphafold3^39^. Visualization and structural alignment were performed using UCSF Chimera and Chimera _X40._

### Microtubule-stimulated ATPase assay

ATPase rates were measured using the EnzChek phosphate assay kit (Molecular Probes, ThermoFisher Scientific). All microtubule-stimulated ATPase assays were performed in assay buffer containing 1XBRB80 with 1 mM ATP, 7 μM taxol-stabilized microtubules. Proteins were used at the following concentrations: 1 μM KIF7-DM or KIF7-DMK_100A_, and with or without 4 μM GLI2-ZF. A reaction mixture containing microtubules and ATP, but no KIF7DM or GLI2-ZF proteins, was used as a blank. The phosphate released upon ATP hydrolysis was monitored at OD360 in a SpectraMax microplate reader every minute for 1 hour. All reactions were initiated by the addition of ATP. The initial reaction velocity was used to determine the ATPase rate. The mean and standard deviation of the ATPase rate from 3 independent experiments was plotted.

### Total Internal Reflection Fluorescence Microscopy Assays

*In vitro* TIRF-based microscopy experiments were carried out as described in Subramanian et al., 2013^41^. Microtubule binding assay: X-rhodamine/HiLyte 647 and biotin-labeled microtubules were polymerized in the presence of GMPCPP, a non-hydrolyzable GTP-analog, and immobilized on a neutravidin-coated glass coverslip. Coverslips were briefly incubated with casein to block non-specific surface binding before the addition of 100nM kinesin in assay buffer and antifade reagent (25mMglucose, 40 mg/ml glucose oxidase, 35 mg/ml catalase, and 0.5% β-mercaptoethanol). Images were acquired from multiple fields of the same chamber.

### Image Analysis

ImageJ was used to extract microtubule-associated GFP fluorescence intensities for TIRF assays. Average intensity per pixel was calculated using rectangular regions-of-interest with a width of 3 pixels and the length of each microtubule. Local background intensities were subtracted using regions of interest of the same area around the selected microtubule. Intensities were not analyzed for microtubules found at the edges of the camera’s field of view. All TIRFM data were analyzed using GraphPad Prism.

### Cell Culture and Immunofluorescence

HeLa cells were obtained from Dr. Robert Kingston (Massachusetts General Hospital). FlpIn-NIH3T3 (ThermoFisher, R76107) cells were purchased, and the Flp-In system was used to overexpress HALO-tagged mouse GLI2 protein. HeLa and NIH3T3 cells were maintained in high-glucose Dulbecco’s modified Eagle’s medium (DMEM) supplemented with 10% fetal bovine serum (FBS), sodium pyruvate (1mM), and L-glutamine (2 mM). Cells were cultured on clean coverslips under the following conditions: 5% CO_2_, 95% air condition at 37^°^C. Cells were grown to ∼70% confluency and transfected with plasmids expressing GFP-tagged KIF7-DM_WT_ or KIF7-DM_K100A_ using jetPRIME transfection reagent and incubated for 18-22 hours. HeLa cells were then fixed using a mixture of methanol and acetone (1:1 in volume) for 10 minutes at −20C, washed with washing buffer (PBS + 0.05% Tween 20), and mounted on 25mmX75mm glass slides with ProLong Diamond Antifade Mountant with DAPI (Thermo Fisher). NIH3T3 cells were subsequently serum-starved with 0.2% FBS in DMEM to induce ciliogenesis for 24 hours, followed by treatment with 500nM SAG for 12-18 hours. Cells were incubated with HALO-JFX-546 dye for 1 hour at 37°C, followed by 3 washes with PBS before fixation for immunofluorescence. Cells were then fixed using a 1:1 v/v mix of methanol and acetone for 10 minutes at −20 °C, washed with washing buffer (PBS + 0.05% Tween 20) 3 times, blocked with blocking buffer (PBS + 2%BSA; OmniPur BSA; EMD Millipore) for one hour at room temperature. Samples were probed overnight at 4°C with the Alexa Fluor 647nm labeled anti-acetylated α-tubulin antibody (Santa Cruz Biotechnology; catalogue# sc-23950; clone# 6-11B-1; 1:500 dilution in blocking buffer). Samples were mounted on a 25mmX75mm glass with ProLong Diamond Antifade Mountant (Thermo Fisher). Z-stacks were acquired on an inverted Nikon confocal microscope using a laser illumination source (488nm, 561nm, 647nm) with a pinhole of 1. Z-projections were generated by sum of images from planes that included the entire cilium.

---

**Extended Data Figure 1.**
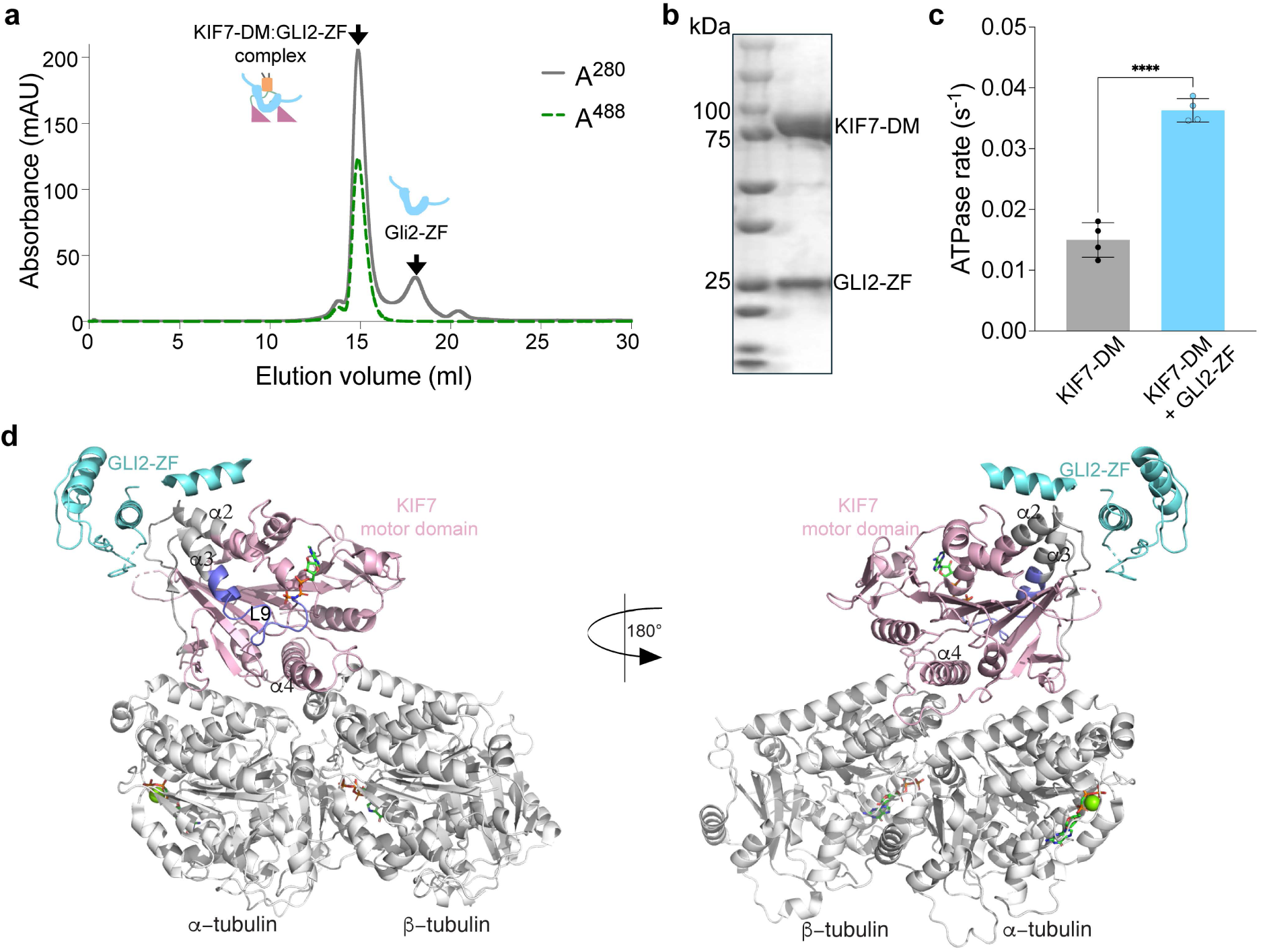
(a) Chromatograms from size exclusion chromatography of a mixture of 125 µM KIF7-DM and 750 µM GLI2-ZF (Superdex 200 10/300 GL) showing the complex that was used for HDX-MS experiments (b) SDS-PAGE of the complex showing bands for KIF7-DM (89kDa) and GLI2-ZF (22kDa) (c) ATPase assay showing microtubule-stimulated ATPase rate of 1 µM KIF7-DM in the absence and presence of 4 µM GLI2-ZF. (d) Cryo-EM structure (PDB ID: <u>7RX0</u>) showing KIF7 motor domain (pink) bound to microtubules (grey) in the presence of GLI2-ZF (cyan) and AMPPNP. The peptides that show protection in HDX-MS upon GLI binding (purple) are not in close proximity to the density corresponding to GLI2-ZF that was observed in the structure.

**Extended Data Figure 2.**
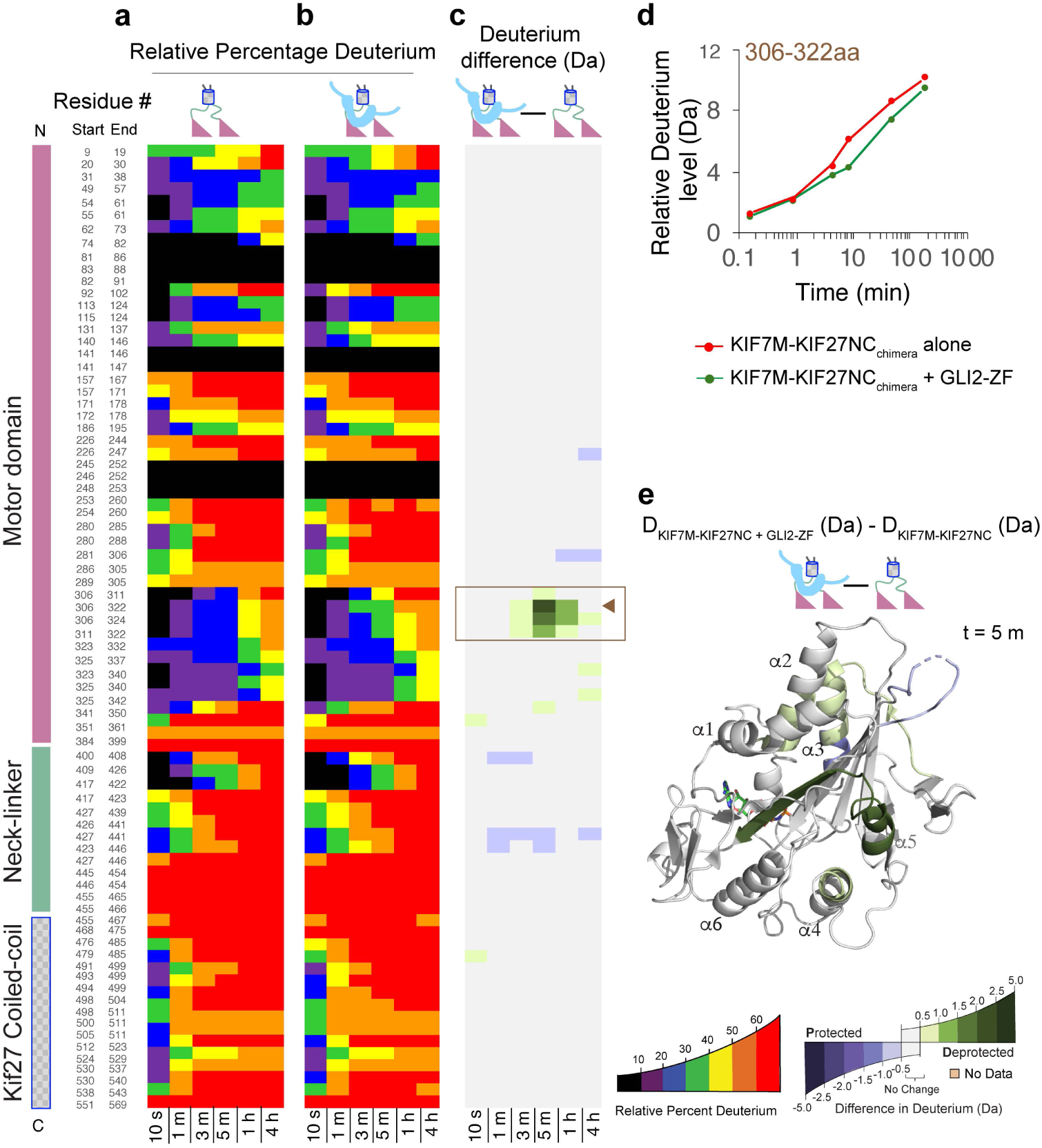
(a) Chiclet plot for relative percentage deuterium for KIF7M-KIF27NC_chimera_. The X-axis shows the time points, and the Y-axis shows each peptide. The relative percentage of deuterium is colored according to the scale shown. (b) Chiclet plot for relative percentage deuterium for KIF7M-KIF27NC_chimera_ in the presence of excess GLI2-ZF is shown. The X-axis shows the time points, and the Y-axis shows each peptide. The relative percentage of deuterium is colored according to the scale shown. (c) Deuterium difference between (a) and (b). The X-axis shows the time points, and the Y-axis shows each peptide. Only the peptides that are coincident in both conditions (KIF7M-KIF27NC_chimera_ and KIF7M-KIF27NC_chimera_ + GLI2-ZF) are shown. The deuterium difference data are calculated by subtracting the absolute deuterium incorporation values for KIF7M-KIF27NC_chimera_ from KIF7M-KIF27NC_chimera_ + GLI2-ZF. The differences are colored according to the scale shown. (d) Deuterium incorporation curve of a selected peptide, which is indicated with solid brown arrowheads in (c). (e) Deuterium difference from 5 m time point in (d) are mapped on KIF7 motor domain (PDB ID: <u>7RX0</u>) with deuterium difference color scale.

**Extended Data Figure 3.**
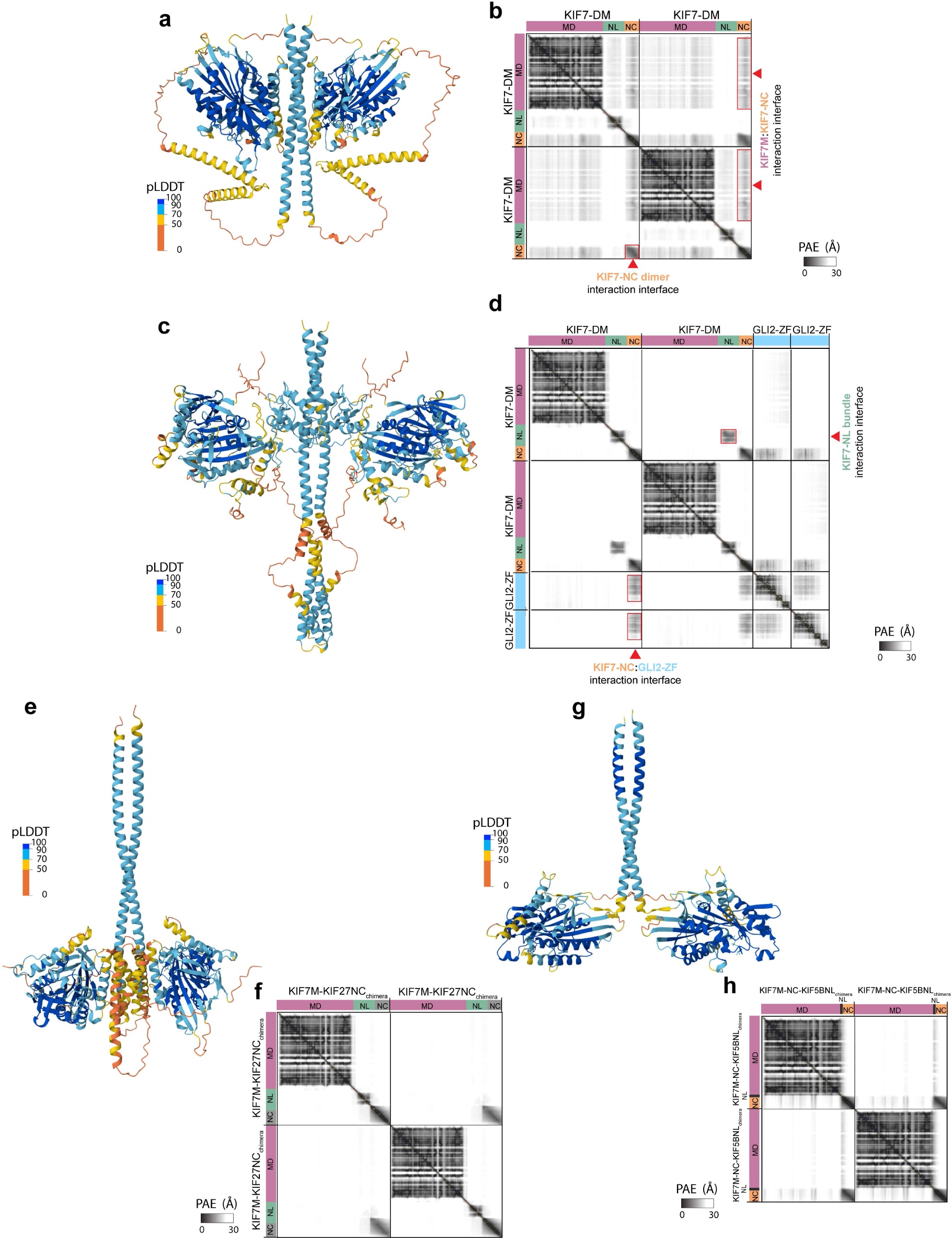
(a) AF3 structural model of KIF7-DM (Figure3a) color-coded by pLDDT score (b) PAE plot for KIF7-DM model in (a) (c) AF3 structural model of KIF7-DM + two copies of GLI2-ZF (Figure3b) color-coded by pLDDT score (d) PAE plot for KIF7-DM:2GLI2-ZF model in (c) (e) AF3 structural model of KIF7M-KIF27NC_chimera_ (Figure3c) color-coded by pLDDT score (f) PAE plot for KIF7M-KIF27NC_chimera_ model in (e) (g) AF3 structural model of KIF7M-KIF5BNL-KIF7NC_chimera_ (Figure3d) color-coded by pLDDT score (h) PAE plot for KIF7M-KIF5BNL-KIF7NC_chimera_ model in (g)

**Extended Data Figure 5.**
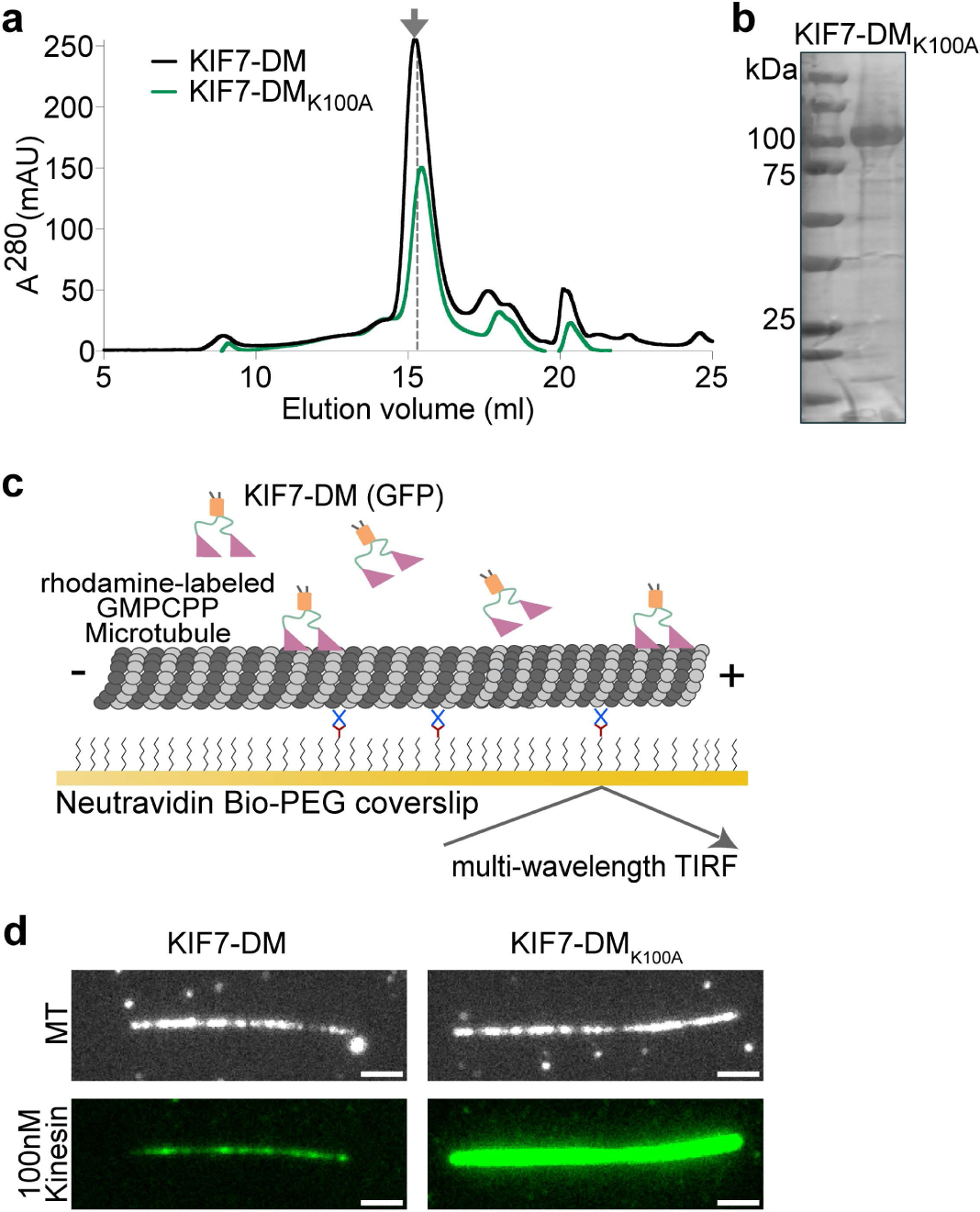
(a) Chromatograms from size exclusion chromatography of KIF7-DM (black) and KIF7-DM_K100A_ (green) on Superdex 200 10/300 GL. Dotted line and arrow indicates elution volume of each protein that was used for experiments (b) SDS-PAGE of purified KIF7-DM_K100A_ (c) Schematic of the *in vitro* total internal reflection fluorescence (TIRF) microscopy-based assay used to examine microtubule-localized KIF7-DM. Rhodamine-labeled GMPCPP-stabilized microtubules (grey) were immobilized on a PEG-treated glass coverslip (yellow) via neutravidin-biotin linkages (blue and brown respectively). GFP-tagged KIF7-DM and 1mM ATP were subsequently added to examine the binding of kinesin on microtubules. (d) Representative fluorescence images of Rhodamine-microtubule (MT) and associated GFP fluorescence of KIF7-DM (left) or KIF7-DM_K100A_ (right). Experiments were performed with 100 nM KIF7 proteins. Fluorescence intensities are displayed using an identical scale for comparison. Scale bars represent 2μm.

